# Post-Stimulus Encoding of Decision Confidence in EEG: Toward a Brain-Computer Interface for Decision Making

**DOI:** 10.1101/2022.11.01.514790

**Authors:** Nitin Sadras, Omid G. Sani, Parima Ahmadipour, Maryam M. Shanechi

**Affiliations:** Ming Hsieh Department of Electrical and Computer Engineering, Viterbi School of Engineering, University of Southern California, Los Angeles, CA, United States of America; Department of Biomedical Engineering, Viterbi School of Engineering, University of Southern California, Los Angeles, CA, United States of America; Department of Computer Science, Viterbi School of Engineering, University of Southern California, Los Angeles, CA, United States of America; Neuroscience Graduate Program, University of Southern California, Los Angeles, CA, United States of America

**Keywords:** EEG, brain-computer interface, confidence, decision-making, source localization

## Abstract

When making decisions, humans can evaluate how likely they are to be correct. If this subjective confidence could be reliably decoded from brain activity, it would be possible to build a brain-computer interface (BCI) that improves decision performance by automatically providing more information to the user if needed based on their confidence. But this possibility depends on whether confidence can be decoded right after stimulus presentation and before the response so that a corrective action can be taken in time. Although prior work has shown that decision confidence is represented in brain signals, it is unclear if the representation is stimulus-locked or response-locked, and whether stimulus-locked pre-response decoding is sufficiently accurate for enabling such a BCI. We investigate the neural correlates of confidence by collecting high-density EEG during a perceptual decision task with realistic stimuli. Importantly, we design our task to include a post-stimulus gap that prevents the confounding of stimulus-locked activity by response-locked activity and vice versa, and then compare with a task without this gap. We perform event-related potential (ERP) and source-localization analyses. Our analyses suggest that the neural correlates of confidence are stimulus-locked, and that an absence of a post-stimulus gap could cause these correlates to incorrectly appear as response-locked. By preventing response-related activity to confound stimulus-locked activity, we then show that confidence can be reliably decoded from single-trial stimulus-locked pre-response EEG alone. We also identify a high-performance classification algorithm by comparing a battery of algorithms. Lastly, we design a simulated BCI framework to show that the EEG classification is accurate enough to build a BCI and that the decoded confidence could be used to improve decision making performance particularly when the task difficulty and cost of errors are high. Our results show feasibility of non-invasive EEG-based BCIs to improve human decision making.

## 1. Introduction

Brain-computer interface (BCI) technology is typically used to restore functionality to injured or impaired patients. While many such BCIs have been developed using invasive neurophysiology modalities [1–9], non-invasive electroencephalography (EEG) based BCIs have also been used to restore functions such as locomotion, motor control, and communication to impaired patients [10–19]. Beyond the treatment of pathological conditions such as motor impairment and neuropsychiatric disorders, there is also growing interest in developing non-invasive BCIs that can improve individual capabilities [20–24].

Prior work on such BCIs includes memory enhancement [25], drowsiness detection [26–28], driving assistance [29], remote robot control [30], cursor control [31], trust evaluation [32], group decision making [33], error detection [34], and attention monitoring [35,36]. Within this class, one important application of interest is to develop BCIs that help subjects make critical decisions with increased reliability, speed, and accuracy, especially in critical, stressful or time-pressured situations [37].

As an example, here we envision a non-invasive confidence-based BCI that aims to improve decision accuracy. To be realized in the future, such a BCI will first need to decode the user’s decision confidence, such that decoded confidence can be used to determine when to provide sensory feedback designed to increase the user’s decision accuracy, as shown in Figure 1. Indeed, self-reported confidence has been shown to be highly predictive of decision accuracy [38–40], and is also reflected in neural activity [39,41,42] and could therefore be an appropriate brain state for use in a BCI for improved decision accuracy. However, developing such a confidence-based BCI for decision making is challenging because it requires answering two questions, which remain elusive.

**Figure 1:**
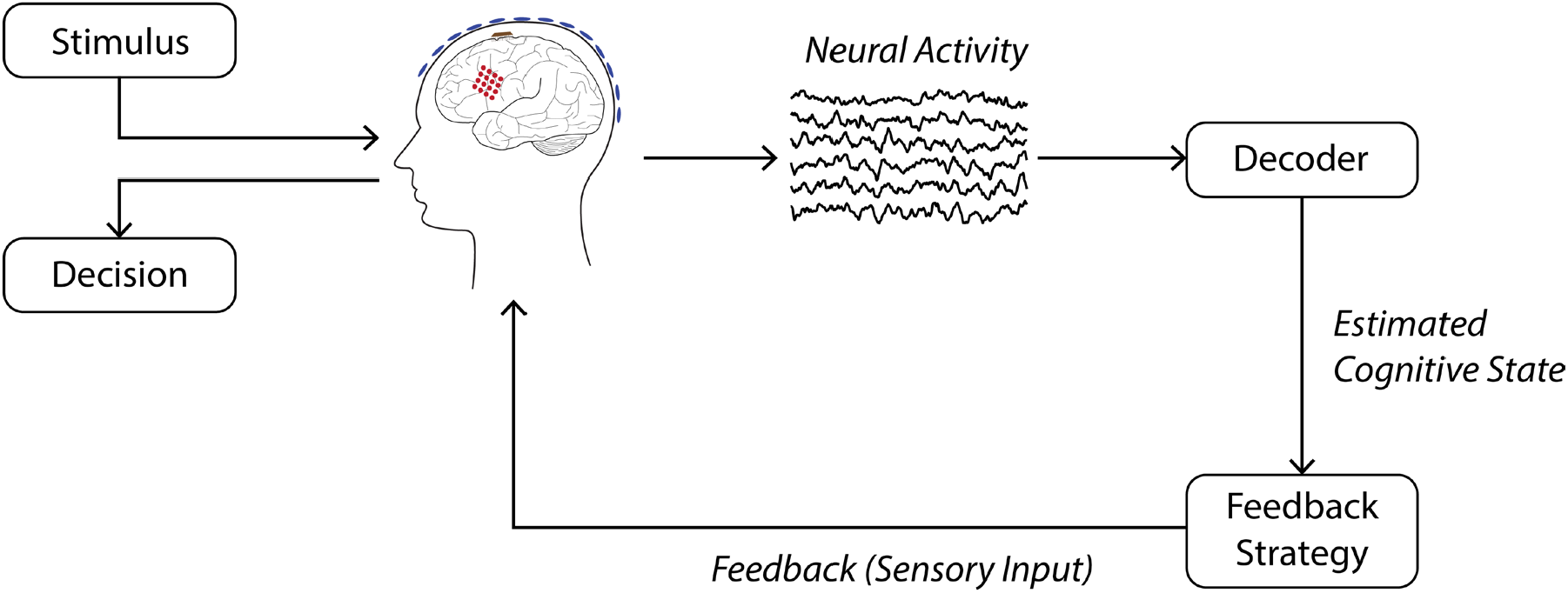
Schematic of a brain-computer interface for improved task performance. Here, the user is performing a task that involves making a decision about a stimulus. Post-stimulus (pre-response) neural activity is passed to the decoder, which estimates the user’s cognitive state (e.g., decision confidence). Based on this estimated cognitive state, feedback can be given to the user in the form of sensory input in order to help improve task performance.

First, we should determine if confidence can be reliably decoded *prior* to the user’s decision, and how accurate such decoding can be. This pre-response decoding is necessary in order for the BCI to have time to influence the user’s decision via sensory feedback. Second, we should determine if the accuracy of such a pre-response decoder is sufficient to enable a BCI that improves the user’s decision accuracy, and under what task conditions this improvement is observed. Here, we address these two standing questions.

Regarding the first question, prior work has shown that decision confidence is reflected in neural activity, and can be decoded at above-chance levels [39,41–44]. However, it is still largely unclear whether the neural encoding of confidence is stimulus-locked or response-locked. This distinction is of critical importance for BCI development, as a BCI can only provide useful information to a user if confidence can be decoded *prior* to the decision – otherwise it is too late to provide such information, as the decision is already made. While some studies show that neural activity is modulated by confidence post-stimulus [41,45,46], others suggest that this encoding is stronger post-response [39,42,47]. We thus first focus on the question of whether confidence is a stimulus- or response-locked phenomenon, and if it is stimulus-locked, how well it can be decoded.

In order to dissociate stimulus-locked and response-locked correlates of confidence, we design a novel experimental task that collects subjective confidence reports while clearly separating the stimulus processing from the response using a time gap. This gap is important as stimulus-locked effects can interfere with response-locked phenomenon without a sufficient post-stimulus gap [48–51]. Results in prior studies differ from each other on both the presence of a post-stimulus gap, and on whether the neural representation of confidence is found to be response-locked or stimulus-locked. For example, in [39] and [42], no post-stimulus gap is present, and confidence appears to be more strongly represented in the response-locked epoch. However, in [41], a post-stimulus gap is used, and it is shown that confidence can be decoded from stimulus-locked neural activity, while the existence of any response-locked confidence modulation is not explored. A study that compares EEG activity collected during tasks *both* with *and* without a stimulus-response gap is necessary in order to reconcile these different conclusions, but no such study has been done so far. Here we design novel experiments and use event-related potential (ERP) analysis to systematically probe the effect of this gap on neural correlates. Our ERP analysis reveals the confidence-related activity to be stimulus-locked, which is promising for the use of confidence in a BCI. We then use EEG source localization analysis to explore which brain regions reflect sources of confidence-related activity.

Having characterized the neural correlates of confidence and their stimulus vs response-locked nature, we then investigate how well confidence can be decoded from single-trial neural activity *prior to* any response. Prior studies have shown that confidence can be decoded from single-trial EEG activity, but it is unclear which of the many existing classification methods is best suited for EEG decoding in this setting. Further, here we can assess the decoding purely due to pre-response activity. This is because the gap introduced in our experiment ensures that the neural activity used for classification does not contain response-related components, which may be influenced by the response modality (verbal, button press, etc.) and which can help/confound the decoding. Thus, we perform a systematic comparison of classification methods for pre-response EEG, including various neural networks, in terms of their pre-response decoding of confidence. Among the considered alternatives, we find support vector machine (SVM) classifiers to have the best performance, likely because of their data-efficient training.

Regarding the second question, even if confidence can be decoded from single trials pre-response, it remains unclear if, and under what conditions, the achieved decoder accuracies can enable a BCI to improve a user’s decision accuracy. We thus next explore this question by devising a simulated BCI framework. Our simulation framework not only allows us to determine whether a BCI can improve task performance, but also under what task conditions such improvement is more pronounced. Our simulation analysis shows that neural classifiers of confidence that have the same accuracy as that observed in our data can indeed improve task performance within the BCI. Further, this improvement is largest for difficult and high-risk task conditions.

## 2. Methods

### 2.1. Experimental task

All experiments were approved by USC’s Institutional Review Board. We developed an experimental paradigm in order to evoke different levels of confidence in response to realistic stimuli. The 3D models and images used in our experiment are similar to those used in [44]. The timeline for a single trial of the experiment is shown in Figure 2A. The stimulus, a character wearing either a cap or a helmet, was shown against a corridor background for 250ms. After a 1.75 s post-stimulus gap, subjects were asked whether they saw a cap or a helmet. This post-stimulus gap was necessary in order to dissociate stimulus- and response-locked phenomena, and is discussed in detail below. Following the post-stimulus gap, subjects reported their response via mouse click – left click for helmet, right click for cap. There was no time limit for this response. After a 1.5 s post-response gap, subjects were then asked to report their confidence on a scale from 1 to 10, with a 1 being a random guess and a 10 being complete confidence. The prompt for confidence report was shown for a fixed duration of 2 seconds, after which the value indicated by the subject was recorded as their confidence. Subjects performed a total of 640 trials in 40-trial blocks. We now describe the rationale behind the gap sizes and the method by which task difficulty and thus confidence level were varied.

**Figure 2:**
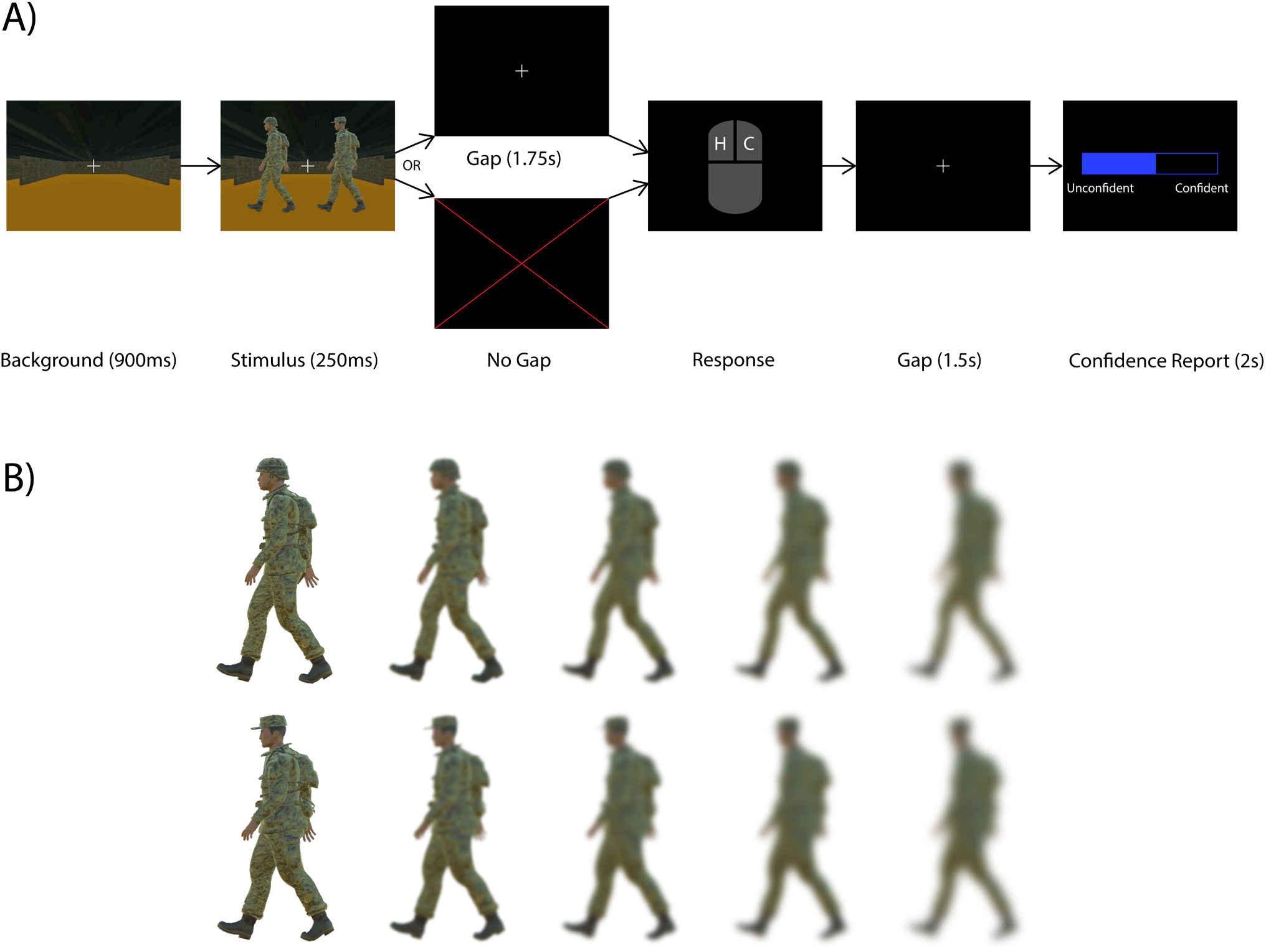
Experimental protocol for the gap / no gap stimulus discrimination tasks. **(A)** Users were presented with an image of a character wearing either a cap or a helmet. In the gap task, a blank screen with a fixation cross was shown for 1.75s before the subject was prompted for a response. In the no-gap task, the subject was prompted for a response immediately. Following a second gap of 1.5s, subjects were then asked to report their confidence via scroll bar on a scale from 0 (unconfident) to 10 (confident). A 0 corresponds to a random guess, while a 10 corresponds to complete certainty. Subjects had a total of 2 seconds to complete their confidence report, but there was no time limit for the initial response. **(B)** Stimuli were blurred to one of five levels to modulate task difficulty. In the first block of trials, all blur levels occurred with equal frequency. In subsequent blocks, the proportion of blurred and unblurred stimuli was adjusted based on the user’s performance to have roughly equal numbers of confident and unconfident trials.

In order to vary the difficulty of the task, we blurred the stimuli by different amounts on each trial. Specifically, the stimuli were blurred to 5 different levels (Figure 2B) using gaussian kernels of varying sizes, with each blur level appearing an equal number of times in the first block. After the first block, the number of maximally- or minimally-blurred stimuli in each subsequent block was adjusted based on the subject’s accuracy in the most recent block. If the accuracy was below 65%, 5 maximally-blurred stimuli were replaced with minimally-blurred stimuli in order to make the task easier. If the accuracy was above 85%, 5 minimally-blurred stimuli were replaced with maximally-blurred stimuli in order to make the task harder. This was done in order to keep the accuracy in the range of 65%-75% because this range results in roughly equal numbers of confident and unconfident trials as follows. For highly confident trials, we would expect near 100% accuracy, while for minimally confident trials, we would expect near 50% accuracy (random chance). Therefore, with an equal number of confident and unconfident trials, the total accuracy should be around 75%. We refer to this task as the *gap task*.

Our 1.75 s post-stimulus gap was necessary in order to separate stimulus- and response-locked phenomenon. ERPs such as the motor-related cortical potential (MRCP) have been shown to start up to 1.5s prior to a response-related movement [52], so it was necessary for the post-stimulus gap to be longer than this. This gap prevents response-locked activity which may be influenced by the response modality (verbal, button press, etc.) from interfering with stimulus-locked activity. Therefore, this gap allows us to observe stimulus-locked activity that is independent of the response modality. Further, without this gap, it is possible that stimulus-related activity can leak into the response-locked epoch, thereby causing stimulus-locked activity to be seen as response-locked. In order to assess the impact of the post-stimulus gap, we also collected data during a stimulus-discrimination task that is identical to the gap task but with no post-stimulus gap, referred to here as the *no-gap task* (Figure 2A).

### 2.2. Data collection

We recruited 16 subjects to participate in our study. 11 subjects participated in the gap task, while 8 subjects participated in the no-gap task. 3 subjects that participated in the no-gap task also participated in the gap task. Two gap-task subjects were removed from the ERP and source localization analyses due to excessive eye movement artifacts. We collected EEG data using a 256 electrode BioSemi ActiveTwo system. In addition to scalp EEG, 2 electrodes were placed on the ears for re-referencing. Further, 4 electrodes were placed around the subjects’ eyes in order to collect electrooculogram (EOG) data. This EOG data was used to correct for eye-movement artifacts, as explained in section 2.3. Stimuli were presented on an 11-inch laptop screen with a 60 Hz refresh rate, using the PsychoPy python toolbox [53]. Subjects were seated approximately 70 cm from the screen, and used a USB-connected mouse for all task-related responses. All subjects were right-handed and controlled the mouse with their dominant hand.

### 2.3. Data pre-processing

We first down-sampled the raw EEG data from 2048 Hz to 256 Hz. We then applied a 1-40 Hz bandpass filter (zero-phase order 1406 Chebyshev FIR filter) to remove DC offsets and high-frequency noise. In order to remove eye-movement artifacts, we performed independent component analysis (ICA) using the InfoMax algorithm [54,55]. We regressed IC’s onto vertical and horizontal EOG signals and removed the IC’s with *r*^2^ values above a threshold of .4. This procedure resulted in 1-3 IC’s being identified for removal. Next, EEG channels that were identified based on a visual inspection as too noisy (mean=13.5, std=11.8 channels removed) were replaced via spatial spline interpolation. Lastly, data was re-referenced to the average of signals from the two ear electrodes. Finally, trials identified based on a visual inspection as containing high noise levels were removed from the analysis. For the gap task, 101 of 6840 trials were removed, and for the no-gap task, 0 of 5120 trials were removed. All preprocessing was done using the FieldTrip toolbox [56].

### 2.4. Data analysis

#### 2.4.1 ERP analysis

We performed an ERP analysis by averaging EEG activity across trials and subjects. We grouped trials by confidence and compared confident trials with unconfident trials during stimulus- and response-locked epochs. Specifically, we performed two analyses. In our first/main analysis, the ‘confident’ group contained trials within the top 20% of confidence reports across recording sessions, while the ‘unconfident’ group contained trials within the bottom 20% of confidence reports across sessions. The remaining trials that had intermediate confidence levels were excluded from both groups due to the ambiguity in the ground truth. For the gap task 3306 out of 6739 trials were excluded. For the no-gap task 2158 out of 5120 trials were excluded. However, in order to ensure that excluding trials did not change our conclusions, we performed a second analysis. This time, we repeated the ERP analysis using all trials, taking the trials with the top 20% of confidence reports across recording sessions as the ‘confident group’, and all the remaining trials in the ‘unconfident’ group. This additional analysis is described in Appendix B and led to consistent conclusions. We performed the ERP analysis for both the gap and no-gap tasks, with the goal of investigating how response- and stimulus-locked correlates of confidence change in the presence of a post-stimulus gap.

To test the statistical significance of the difference in ERP between confident and unconfident trials, we used a cluster-based permutation test [57]. This method has been shown to solve the multiple-comparisons problem, which arises when many statistical tests are performed simultaneously. The procedure for the cluster-based permutation test is explained in Appendix A.

#### 2.4.2 EEG source localization

To understand the neural sources of the confidence-related activity seen in our ERP analysis, we performed source localization. Indeed, the high-density 256 channel EEG system makes our experiments well-suited for this localization compared to experiments with lower channel counts [58,59]. EEG activity measured on the scalp is caused by intracranial electrical currents. Distributed source localization algorithms attempt to solve the inverse problem of estimating the 3D distribution of intracranial current density from scalp EEG activity [60–62]. One such algorithm, eLORETA, has been shown to solve this inverse problem with exact, zero-error localization even in the presence of noise [63]. Further, several studies have shown that sources identified by the LORETA family of algorithms are consistent with those identified by neuroimaging methods such as MRI [59,64–68]. It has also been shown that EEG systems with high electrode density, such as the one used in our study, improve localization performance [58,59]. In order to localize the sources of confidence-related EEG activity, we used the eLORETA algorithm as implemented in the FieldTrip MATLAB toolbox [56]. The eLORETA algorithm requires a volume conduction model of the head (head model) to determine how electrical activity generated at the source points propagates to the scalp. We used a template head model provided by the FieldTrip toolbox that is described in [69]. We also performed a control analysis in Appendix C and Figure A2 using a public dataset with simultaneous EEG and intracranial stimulation, with the stimulation electrode positions serving as the ground truth for source localization analysis. Using this control analysis, we validated our source localization pipeline to show its accuracy and further found that performance when using a subject-specific head model did not significantly differ from performance when using a template head model.

### 2.5 Single-trial decoding

We compared six classification algorithms in terms of their ability to decode confidence from single trials of the gap task (Figure 3). Specifically, we used logistic regression (log), support-vector machine (SVM), a Riemannian geometry (RG) classifier [70,71], and three neural network (NN) architectures. The neural networks included a multi-layer perceptron (MLP) network, a spatial convolutional NN (CNN), and EEGNet, an NN that was designed specifically for use with EEG data [72].

**Figure 3:**
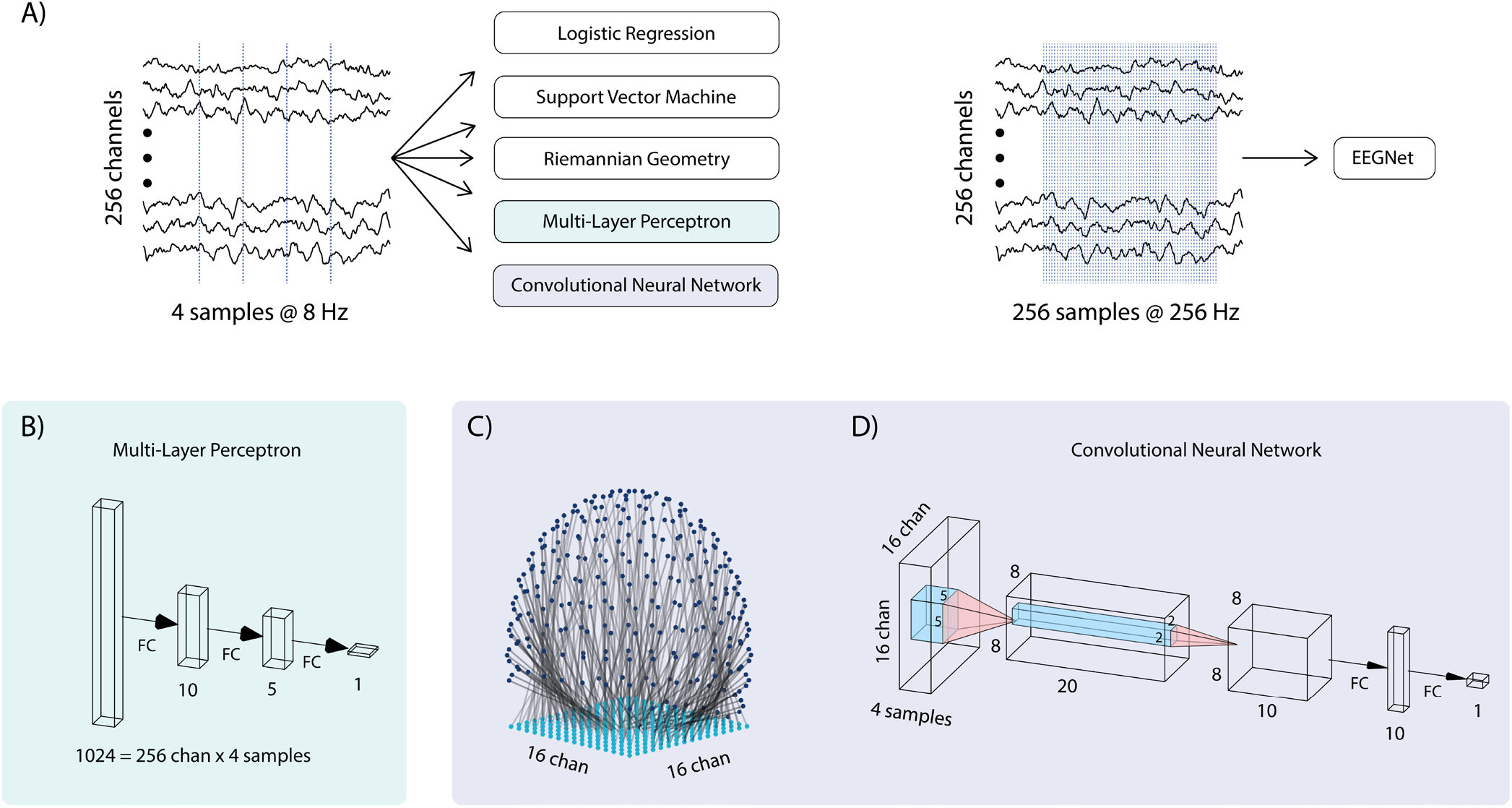
Confidence classifier architecture details. (**A**) 8 Hz data from .2 to .7 s post-stimulus was used as input to the logistic, support vector machine, Riemannian geometry, multi-layer perceptron (MLP), and convolutional neural network (CNN) classifiers. For the EEGNet classifier, we used 256 Hz data from 0-1 seconds post-stimulus. We used a different sampling rate and longer time duration for the EEGNet input in order to match the characteristics of the data used to validate it in prior work. Indeed, more samples are required to get the most out of EEGNet’s temporal convolution layers (despite this, EEGNet was not the best performant classifier, see results). An anti-aliasing filter was applied before downsampling. (**B**) MLP architecture. Blocks indicate data shape at each layer. Four samples from all 256 channels were flattened into a vector of length 1024 before being fed through fully-connected (FC) layers. In both panel B and panel D, the numbers beneath FC layers indicate the number of units in that layer. For the MLP network, the first hidden layer has 10 units, the second hidden layer has 5 units, and the output layer has 1 unit. (**C**) EEG electrodes were mapped onto a 16×16 grid for use with the CNN. Each grid point was mapped to the electrode nearest to it. Since 4 time-samples were used (panel A, left), the resulting tensor was of size 16×16×4. (**D**) CNN architecture. Input data was fed through 2 convolutional layers followed by a single hidden FC layer.

For SVM classification, we used a nonlinear SVM with a Gaussian kernel. The SVM kernel size parameter and the loss regularization parameter were selected with a 2D grid search based on the training data. SVM and logistic regression classifiers were implemented using the scikit-learn Python library [73].

RG classifiers have been shown to be effective in various EEG classification tasks [70,71,74]. In this work, we use a similar approach to that used in [72]. This approach involves dimensionality reduction via xDAWN spatial filtering [75], a nonlinear transformation based on Riemannian geometry [70,71], and classification via logistic regression.

NN classifiers have been shown to have better classification performance than traditional methods in several domains, and have gained popularity for their ability to approximate arbitrary nonlinear functions [76–79]. CNNs are a modification to standard NNs that use 2D convolution to take advantage of spatial structure in input, thereby reducing the number of trainable parameters. CNNs have shown great success in tasks such as image classification, where the input has spatial structure [80–82]. Lastly, EEGNet uses 1D temporal convolutions and has been shown to outperform other classification methods on several EEG datasets [72]. The architectures for our MLP and CNN are shown in Figure 3A.

The input to our spatial CNN is a three-dimensional tensor, where the first two dimensions are spatial, i.e., corresponding to EEG electrode location, and the third dimension is time. In order to use this spatial CNN with our EEG data, we needed a mapping from our 256 three-dimensional electrode locations onto pixels in a two-dimensional 16 by 16 image, which we took as the first two dimensions of the CNN input. It was important that this mapping preserve the spatial structure of the electrode coordinates, i.e., that electrodes that were near each other on the EEG cap were also near each other in the 16 by 16 image. Our method for doing this is illustrated in Figure 3C. Essentially, we placed a 16 × 16 point grid under the 3-D electrode coordinates, and assigned each grid point to the electrode nearest to it. In order to prevent multiple electrodes from mapping to the same grid point, grid points were assigned sequentially, with each grid-point being assigned to the nearest *un-assigned* electrode. The architectures of the MLP and CNN classifiers are shown in Figure 3B and Figure 3D, respectively. A rectified linear unit (ReLU) activation was used as the activation in all hidden units in both MLP and CNN classifiers, and both classifiers were implemented using the tensorflow library [83].

For the EEGNet classifier, we used 256 Hz data from 0-1s post-stimulus as input (Figure 3A, right). For all other classifiers, we used 8 Hz data from 0.2-0.7s post-stimulus as input (Figure 3A, left). We used a different sampling rate and longer time duration for the EEGNet input in order to match the characteristics of the data used to validate it in prior work. Indeed, more samples are required to get the most out of EEGNet’s temporal convolution layers. Note that despite this, EEGNet was not the best performant classifier (see results). To obtain ground-truth binary class labels, we thresholded confidence reports at the 80^th^ percentile across all subjects – trials with a reported confidence at or above this value had a ground-truth label of ‘confident’, while those below this value had a ground-truth label of ‘unconfident’.

We assessed the performance of all the classifiers via the area under the curve (AUC) measure [84]. Briefly, each classifier takes a single trial of neural activity as input, and computes a continuous-valued score as output. This score is then thresholded to determine the predicted discrete class (confident or unconfident) of the associated input. By varying this classification threshold, we also vary the true-positive rate (TPR) and false-positive rate (FPR) of the classifier. The receiver operating characteristic (ROC) curve plots the TPR of a classifier against its FPR for various classification thresholds, and the AUC is the area under this curve. An AUC of 1 indicates perfect classification, while an AUC of .5 indicates that a classifier does no better than chance.

All classifiers were evaluated via 5-fold stratified cross-validation. This was done using the scikit-learn implementation of cross-validation [73].

### 2.6 BCI simulation framework

In addition to studying the neural correlates of confidence and their decoding, our next goal was to determine if confidence decoders with the same accuracy as that observed in our data could be used as part of a BCI to improve a user’s task performance. To do this, we devised a simulated BCI framework based on behavioral data collected in our experimental task. This behavioral data includes decision accuracies and confidence reports. We simulate a general stimulus-discrimination task where a user must determine if a stimulus belongs to one of two categories. The simulated BCI aims to help the user with making the correct decision. To do so, the BCI decodes the user’s confidence after each stimulus presentation and repeats the stimulus until the decoded confidence is above a given threshold. Once this confidence threshold is reached, the user is allowed to respond. Our simulation accounted for the fact that the decoder is imperfect by using the decoding accuracy observed in our EEG data. Our simulation also accounted for the fact that the user’s response accuracy is a function of their confidence based on the behavioral data. Details of how the simulation framework was developed are provided in the results section (section 3.4).

## 3. Results

### 3.1 Confidence-related ERPs are stimulus-locked

We performed a grand-averaged ERP analysis, comparing confident trials with unconfident trials for the gap and no-gap task, for both stimulus- and response-locked epochs (Figure 4A). In the no-gap task, we found that confident and unconfident ERPs at channel Pz were significantly different (p < .05, cluster-based permutation test) both around 500ms after the stimulus onset and around the response time (Figure 4A, top row), thus making it ambiguous whether the difference was stimulus-locked or response-locked. Topographies depicting ERPs across all channels at both these times also had similar characteristics, with positive differences between conditions appearing over the parietal regions (Figure 4B, topography plots 1 and 2). This pattern of activity is consistent with the P300 ERP, which is known to be associated with task-related stimulus processing [85,86].

**Figure 4:**
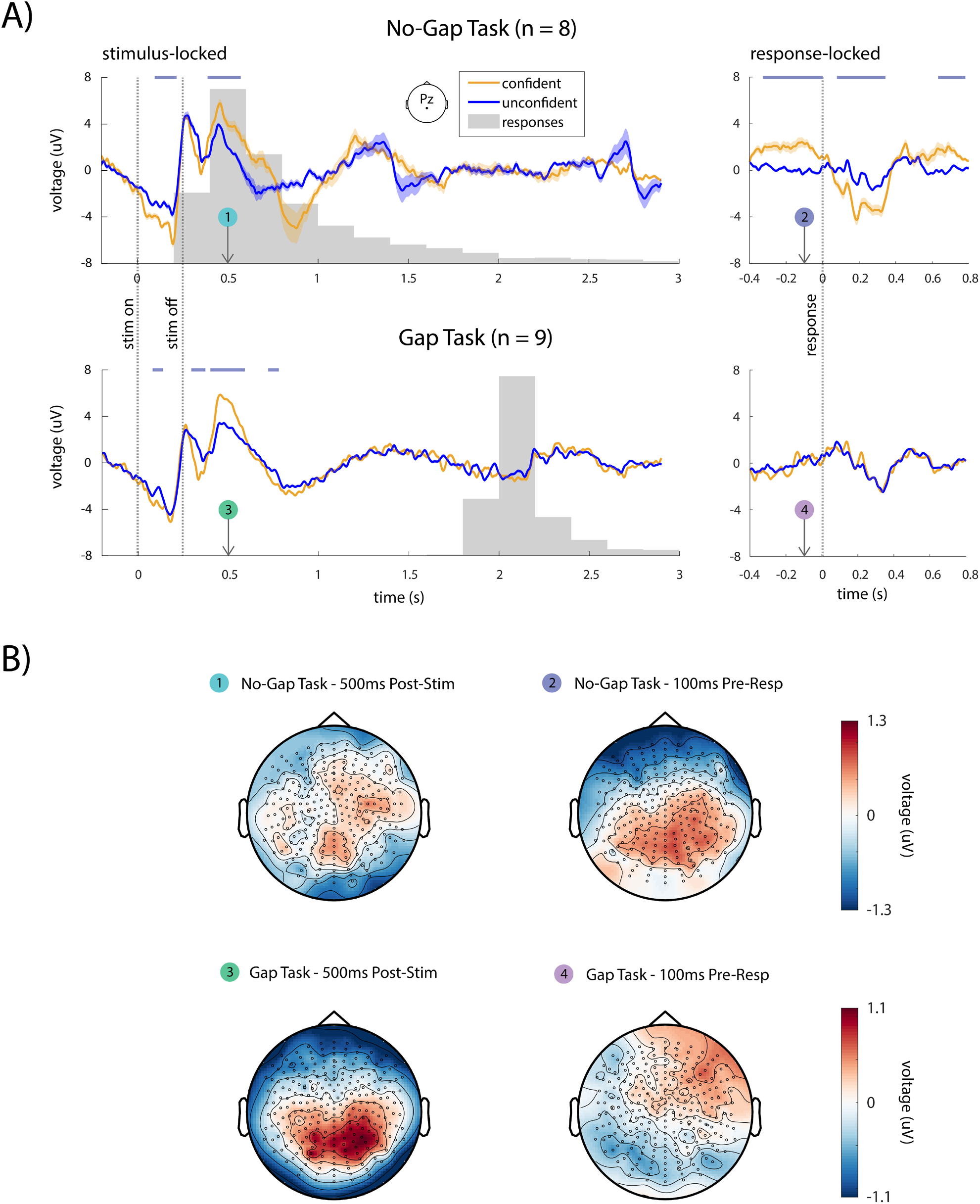
EEG analysis results. (**A**) Grand-average stimulus-locked (left panels) and response-locked (right panels) ERPs at electrode Pz for confident and unconfident trials for the no-gap task (top panels, n=8 subjects) and gap task (bottom panels, n=9 subjects). Horizontal lavender lines above each plot denote regions where ERPs are significantly different (p < .05, cluster-based permutation test). (**B**) Topographic plots are shown for the difference between confident and unconfident conditions at 500 ms post-stimulus (left panels, plots 1 and 3) and 100 ms pre-response (right panels, plots 2 and 4). In the no-gap task, differences between confident and unconfident conditions can be seen over parietal channels in both the stimulus-locked epoch (plot 1) and the response-locked epoch (plot 2). In the gap task, a clear difference is seen in the stimulus locked epoch, but not in the response-locked epoch. Further, the topographies in the response-locked epoch of the no-gap task (plot 2) and the stimulus-locked epoch of the gap task (plot 3) have similar characteristics. These results suggest that the response-locked confidence-related activity in the no-gap task is actually a stimulus-locked phenomenon that is confounded by the lack of a gap.

Interestingly, unlike the no-gap task in which there was ambiguity as to whether the activity was stimulus- or response-locked, in the gap task, this pattern of activity was only observed in the stimulus-locked epoch, and not in the response-locked epoch (Figure 4A, bottom row; Figure 4B, topography plots 3 and 4). Importantly, in the gap task, this stimulus-locked difference occurred long before the response, suggesting that there is no possibility of interference between stimulus- and response-locked activity.

Parietal regions were among the regions with the largest ERP differences for the stimulus-locked ERP in the no-gap task, the response-locked ERP in the no-gap task, and the stimulus-locked ERP in the gap task, suggesting that shared brain processes may be involved in all three epochs (Figure 4B, topography plots 1-3). Given that the response-locked differences in ERP follow a different pattern in presence of a post-stimulus gap (Figure 4B, topography plot 4), these results suggest that confidence-related activity in our task is stimulus-locked and not response-locked. These results suggest that in the absence of a gap, it is the stimulus-locked activity that appears as response-locked. When a gap is added, stimulus-locked and response-locked activity are separated, thus removing the ambiguity.

### 3.2 Cortical sources of confidence-related activity

Having identified that confidence-related activity is stimulus-locked in the presence of a gap, we then used eLORETA source-localization analysis to explore which brain regions contain sources of confidence-related activity in the gap task. We used a template head model and template electrode coordinates in order to perform this analysis (section 2.4.2). We first confirmed that using a template head model gives sources that are comparable to those obtained using subject-specific head models based on MRI data and co-registered EEG electrode locations. To this end, we used the Localize-MI dataset, a publicly available dataset developed for the express purpose of validating source-localization methods [65]. The dataset features high-density EEG data that was recorded while subjects received intracranial electrical stimulation. The locations of the stimulating electrodes serve as the ground-truth for source-localization methods. Our analysis revealed that eLORETA with both subject-specific and template anatomy performed significantly better than chance level in localizing the stimulating electrodes (p ≤ 1.2e-12, paired t-test). Further, performance when using subject anatomy was not significantly better than when using template anatomy (p = .1, paired t-test). This result indicates that template anatomy can indeed be used to perform reliable source analysis. The details of this validation analysis are shown in Appendix C.

Having validated our source analysis pipeline, we now discuss the results of our source analysis on the gap task EEG data. EEG data was first down-sampled to 20 Hz and separated into confident and unconfident trials. Next, we computed a confident trial average and an unconfident trial average for each subject. We then used eLORETA to compute the neural sources for both the stimulus-locked epoch (500ms post-stimulus) and the response-locked epoch (100ms pre-response) for each subject and condition (confident, unconfident). This left us with 1 confident source map and one unconfident source map for each subject. Lastly, we used a cluster-based permutation test across subjects to test if there was a significant difference between confident and unconfident sources. We found a significant difference (p < .01) in the stimulus-locked epoch of the gap task, but not in the response locked epoch. This result supports our previous finding that, in the presence of a post-stimulus gap, confidence-related activity is stimulus-locked and not response-locked. The p-values of clusters below the significance threshold in the stimulus locked epoch of the gap task are shown plotted on a template brain in Figure 5.

**Figure 5:**
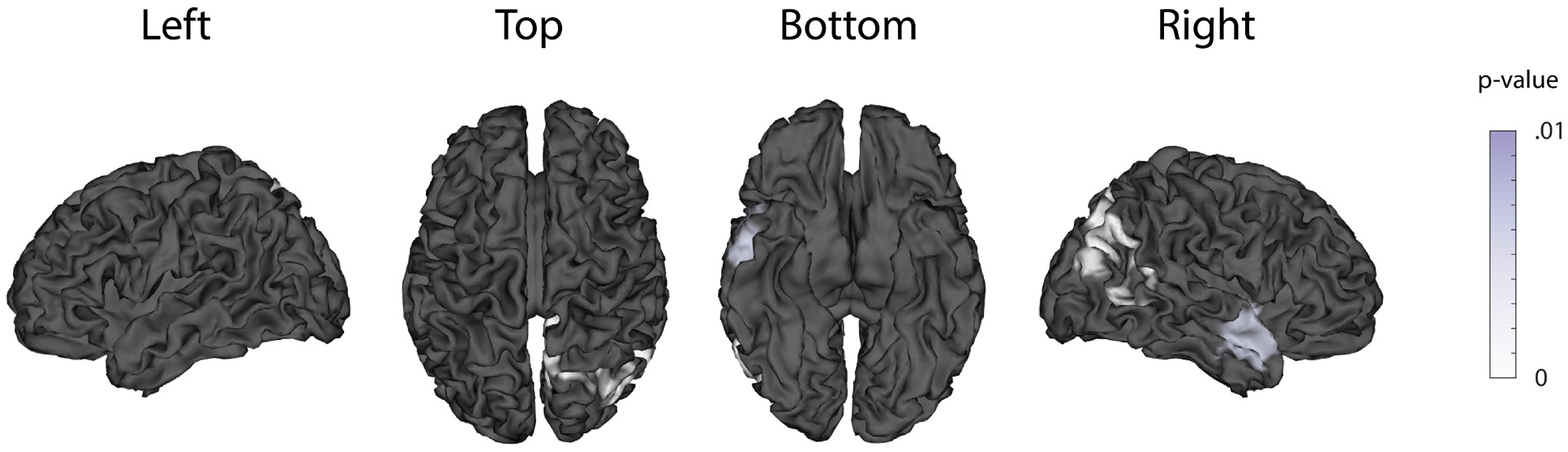
EEG source localization analysis. Neural sources of confidence-related activity 500ms post-stimulus in the gap task, computed using the ELORETA source localization algorithm. A cluster-based permutation test revealed a significant difference between confident and unconfident sources (p < .01). The p-values of below-threshold clusters are shown superimposed on the MNI template brain. No significant difference was found in the response-locked epoch of the gap task.

Our source analysis found below-threshold clusters in the right occipital and temporal lobes, suggesting that these brain regions are involved in the neural representation of confidence. The temporal and occipital lobes are a part of the ventral visual pathway, which is known to be involved in object recognition and visual memory [87]. Our results suggest that these regions may also be involved in representing confidence following decisions in a visual object recognition task. Several types of visual processing have also been shown to be lateralized – for example, evidence suggests that language processing occurs primarily in the left hemisphere, while face recognition occurs primarily in the right hemisphere [88–90]. Some aspects of object-based attention and categorization may also be lateralized to the right hemisphere [91]. These phenomena may contribute to our observation that neural sources of confidence in our task appear to be right-side lateralized.

### 3.3 Confidence can be decoded from single trial stimulus-locked pre-response EEG activity

Our results thus far suggest that the neural correlates of confidence are stimulus-locked, and that these correlates appear prior to the subject’s response when a gap is present. We next sought to decode confidence from single trial stimulus-locked data in the gap task. Note that while various studies have looked at decoding of confidence, here by doing the decoding in the new gap task, we could ensure that the decoding only used the pre-response activity and that the response-related activity was not helping the decoding results. Thus, our decoding can inform whether a BCI that requires reliable decoding pre-response can be constructed.

We used a battery of classifiers, described in section 2.5, to decode confidence from single-trial, stimulus-locked data from the gap task. All classifiers were evaluated via 5-fold stratified cross-validation. Figure 6 shows the average AUC across all subjects (panels A,B) and across folds for each subject (panel C). P-values for across-subject and per-subject results were corrected for multiple comparisons via the Benjamini-Hochberg procedure [92]. For pooled results across subjects, all classifiers were able to classify confidence with significantly above chance accuracy (p ≤ 1e-17, N=55, paired t-test; Figure 6B), suggesting that confidence is robustly decodable from single trial stimulus-locked pre-response EEG activity. This shows feasibility of a BCI that needs to decode prior to response. Further, for 9 out of 11 individual subjects, all 5 classifiers achieved significant decoding, while for the remaining 2 subjects, 3 out of 5 classifiers were significant (p ≤ .05, N=5, paired t-test; Figure 6C). Notably, of all the classification methods, the SVM performed best, with an average AUC of .76 across all subjects and folds (Figure 6B). Also, SVM classification was significant in every one of the 11 individual subjects (Figure 6C). It is likely that the SVM performed better than the more sophisticated deep learning methods because SVM requires relatively fewer training samples for accurate model fitting.

**Figure 6:**
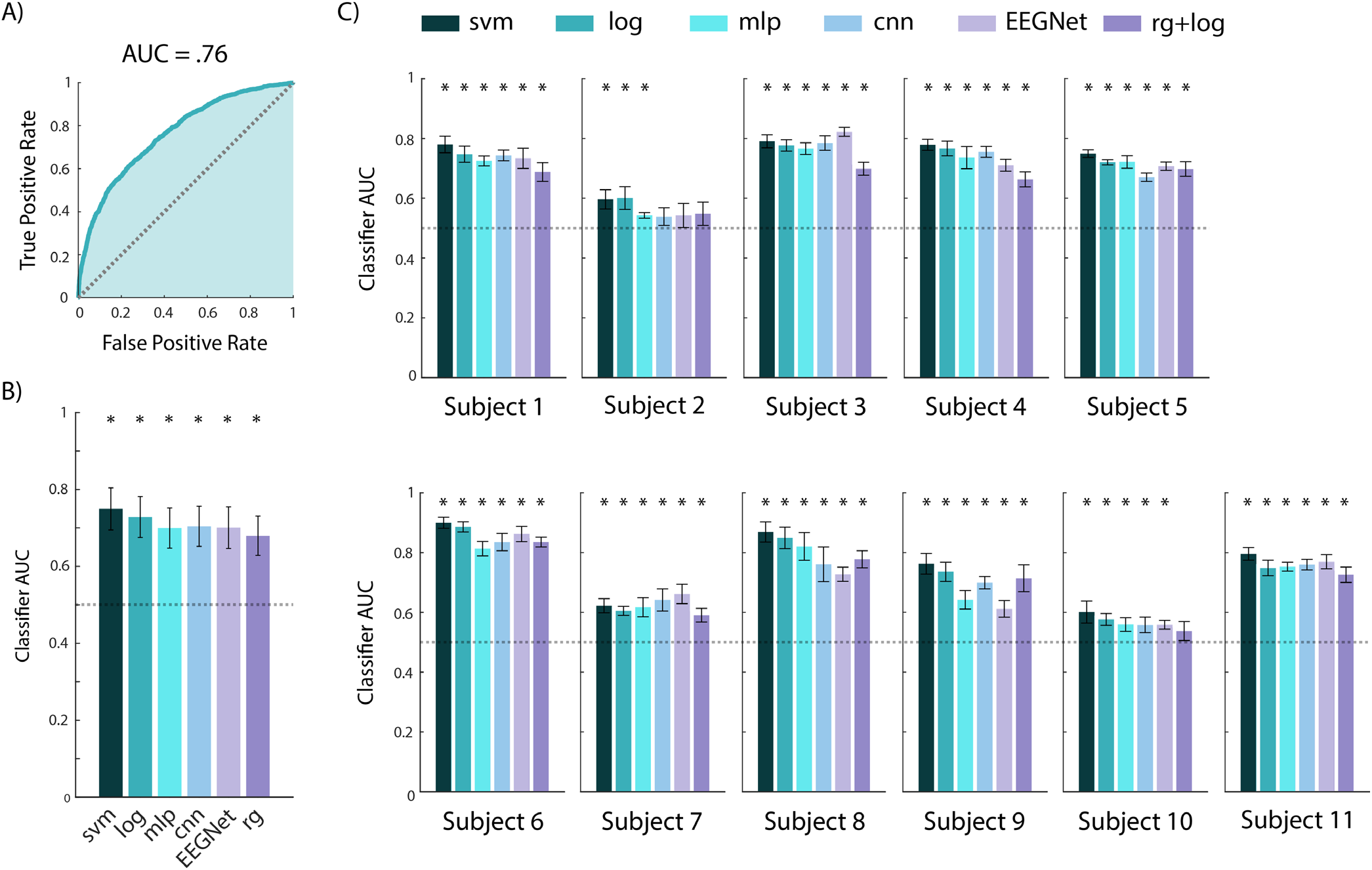
Confidence classification results using just stimulus-locked pre-response activity in the gap task. (**A**) The receiver operating characteristic (ROC) curve for the support vector machine (SVM) classifier, constructed by pooling test trials across all subjects. (**B**) Average performance across all 11 subjects and 5 cross-validation folds, for 6 classification methods. Error bars show the standard deviation. All methods performed significantly above chance level (p < 1e-17). The SVM performed significantly better than the other 5 methods (p < 1e-4). We note that the SVM outperformed EEGNet despite using fewer samples as input. Classifier abbreviations are as follows: svm (support vector machine), log (logistic regression), mlp (multilayer perceptron neural network), cnn (convolutional neural network), rg (Riemannian geometry). (**C**) Classifier performance for each subject, averaged across folds. For all subjects, at least 3 of the 5 classifiers could decode confidence significantly above chance level (p < .05). For 9 out of the 11 subjects, all 5 classifiers could decode significantly above chance (p < .05). Significant decoding (p < .05) is marked with a star. All p-values represent the results of a Benjamini-Hochberg corrected t-test.

### 3.4 Developing a simulation framework for BCIs for improved decision accuracy

As previously discussed in section 2.6, we developed a simulation framework to show how our confidence classifier can be used as part of a BCI to improve a user’s task performance (Figure 7). The details of the developed simulation framework are as follows: when the stimulus is first presented, the simulated user’s true confidence, *c*, is drawn from a distribution based on real confidence reports collected during the gap task. The simulated decoder will then determine if the user’s confidence is above threshold (confident) or below threshold (unconfident). In order to simulate a realistic, imperfect decoder, we set the false-positive and true-positive rates (FPR and TPR) of the simulated decoder to be the same as what the SVM classifier (section 2.5) achieves for confidence classification on the real EEG data, i.e., we use the SVM’s ROC curve in Figure 6A. Specifically, we chose the point on the SVM’s ROC curve that maximizes decoder accuracy when 20% of trials are confident and 80% are unconfident – this reflects the fact that our threshold for confidence is placed at the 80^th^ percentile of the initial confidence distribution. We found this maximum accuracy point on the ROC curve as follows: for a given number of positive (confident) samples *p* and negative (unconfident) samples *n*, the point on the ROC curve with highest accuracy is the point with a tangent line of slope *n/p* closest to the top left corner [93].

**Figure 7:**
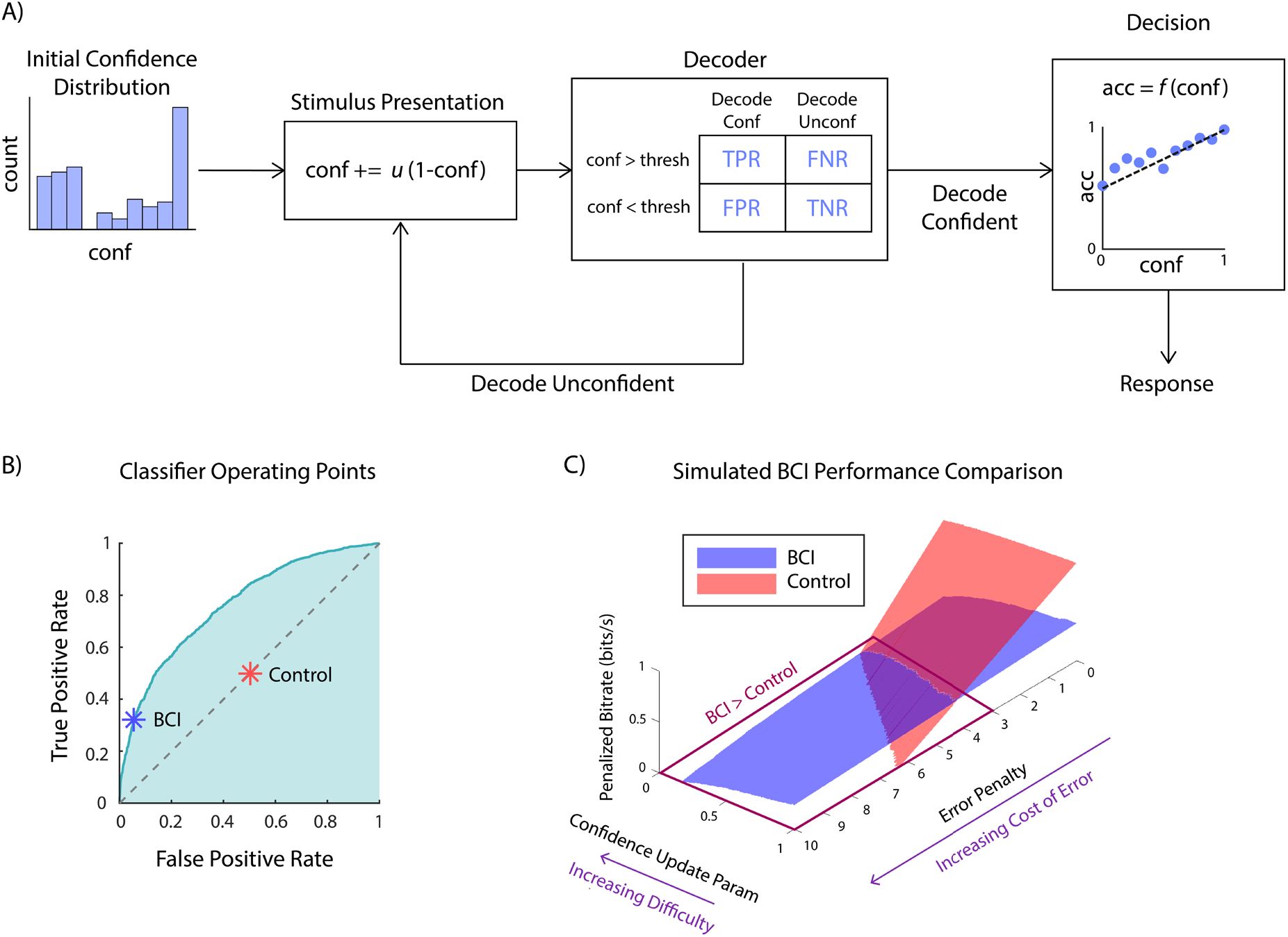
BCI simulation framework showing that confidence classifiers could be used in a BCI to improve decision making. (**A**) Simulation flowchart. First, an initial confidence value is drawn from a distribution determined by subjects’ reported confidence in the collected data. Upon stimulus presentation, confidence is updated based on u, the confidence update parameter. Based on this updated confidence value, we probabilistically simulated whether the decoder classifies the subject as confident or unconfident. We simulate the decoder decision based on the confusion matrix we obtained from real EEG data. The confusion matrix contains the probability of each decoder outcome (top) given the user’s true confidence (left), that is, the true positive rate (TPR), false negative rate (FNR), false positive rate (FPR) and true negative rate (TNR) of the decoder on real EEG data. If the decoder outputs ‘unconfident’, then the stimulus is repeated. If the decoder outputs ‘confident’, the subject is allowed to respond. The subject’s response is simulated as a Bernoulli random variable, with the probability of correctness being a linear function (denoted by f(.)) of their final confidence value. This confidence-accuracy function is described in section 3.4 and is based on the actual behavioral data. (**B**) Operating points for the BCI and control classifiers, shown over the receiver operating characteristic (ROC) curve of the SVM applied to real EEG data. The BCI operating point is the point on the ROC curve that maximizes decoder accuracy when 20% of trials are confident and 80% are unconfident. The control classifier operating point corresponds to declaring confident or unconfident randomly with 50% probability. (**C**) Simulation results. Penalized bitrate for both BCI and Control conditions is plotted against the confidence update parameter and error penalty. For each parameter combination, 100,000 trials were simulated. The results show that the BCI outperforms the control condition in regimes where the confidence update parameter is low (high difficulty) and the error penalty is high (high cost of error).

Given the FPR and TPR corresponding to the chosen point on the ROC curve, the decoder output is simulated as follows. If the user’s confidence is above threshold (confident), then the decoder’s output is drawn from a Bernoulli distribution with parameter *p* equal to the TPR – because TPR is the probability that the decoder will correctly output ‘confident’ when the subject is actually confident. If the user’s confidence is below threshold (unconfident), then the decoder’s output is drawn from a Bernoulli distribution with parameter *p* equal to the true-negative rate (TNR), which is equal to 1 minus the FPR – because TNR is the probability that the decoder will correctly output ‘unconfident’ when the subject is actually unconfident. For both cases, a Bernoulli outcome of ‘1’ corresponds to a decoder output of ‘confident’ while an outcome of ‘0’ corresponds to a decoder output of ‘unconfident’

If the decoder outputs ‘unconfident’, then the BCI repeats the stimulus. We then update the confidence according to the following rule:

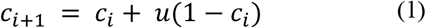

where *c*_*i*_ ∈ [0,1] is the user’s confidence after the *i*th stimulus repetition, and *u* ∈ [0,1] is a confidence update parameter that determines how much the confidence increases after each stimulus presentation. Low values of *u* indicate a difficult task where several repeated presentations of the stimulus are required to gain above-threshold confidence, while high values indicate an easier task where fewer stimulus repetitions are needed. The BCI continues the stimulus repetitions until the decoder outputs ‘confident’, at which point the user is allowed to respond. The user’s decision is simulated by a weighted coin flip, where the probability of correctness linearly increases from .5 at *c* = 0 to .98 at *c* = 1. This linear increase in confidence closely mirrors the confidence-accuracy curve observed in our gap task behavioral data (Figure 7A, ‘decision’ box).

As a control, we also simulated a task where the stimulus was repeated randomly instead of based on a BCI decoder. For the control simulation, after each stimulus presentation, there was a 50% chance that the user would be allowed to respond, and a 50% chance that the stimulus would be repeated. Importantly, to reflect the fact that a post-stimulus gap is needed to decode confidence but not needed for this random control BCI, we designed each repetition of the control task to take less time than each repetition of the BCI task. Specifically, each BCI repetition takes 700 ms, while each control repetition takes 450 ms. The BCI repetition time was chosen because our SVM classifier uses data from up to 700 ms after stimulus onset. The control repetition time was chosen based on a stimulus duration of 250ms and a reaction time of 200ms, for a total of 450ms. This is roughly the amount of time that a subject would need to complete a trial without any BCI intervention.

Task performance within the simulation was measured via a penalized bitrate, which we define as

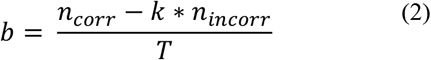

where *n*_*corr*_ is the number of correct responses, *n*_*incorr*_ is the number of incorrect responses, *T* is the total time taken to complete all trials, and *k* ∈ 0, ∞) is the error penalty. Large values of *k* indicate that errors are very costly, while small values indicate that they are not. This penalized bitrate measures the number of correct answers per unit time, but with an additional term that serves to penalize incorrect responses. By varying the parameters *u* (1) and *k* (2), we can simulate tasks with varying levels of difficulty and error cost. We varied *u* from .01 to 1 in steps of .01 and varied *k* from 0 to 10 in steps of 1. For each combination of parameters, we simulated 100,000 BCI trials and 100,000 control trials.

### 3.5 Decoded confidence can be used to improve task performance in a simulated BCI

Having designed the simulation framework for the BCI, we explored the advantage of such a BCI as a function of task difficulty and error cost, which are quantified by *u* and *k* in equations (1) and (2) respectively. In Figure 7D, we plot the penalized bitrate against the parameters *u* and *k* for both the BCI and control cases. In the BCI case, a decoder with the same accuracy as the SVM on real data is used to determine if the subject is confident or unconfident. In the control case, the decoder will output “confident” with a 50% chance, and “unconfident” with a 50% chance, independent of the user’s true confidence. We show only regions of the plot where the penalized bitrate is positive. The qualitative interpretation of a negative bitrate is that the user is making so many errors that they are better off not doing the task at all and having a bitrate of 0. As such, comparisons between BCI and control conditions for negative bitrates are not meaningful.

Our results show that the BCI outperforms the control in the high-difficulty (low *u*), high-error cost (high *k*) regime. In the low-difficulty, low-error cost regime, however, the control has a higher penalized bitrate. This is because control trials are designed to be shorter than BCI trials (section 2.6), and thus for low values of the error penalty *k*, this difference in trial duration outweighs the better decision accuracy that the BCI enables – that is, it outweighs the difference in the bitrate numerator *n*_*corr*_ − *k* ∗ *n*_*incorr*_. In other words, for low values of error penalty *k*, it is better to favor speed over accuracy, and the faster control trials have a higher bitrate. For high error penalties and high task difficulty, this is no longer the case; the longer trial duration of the BCI is offset by the fact that the BCI prevents the user from making costly errors. In other words, when the error penalty *k* is high, it is better to favor accuracy over speed, and thus BCI trials, which are more accurate, achieve a higher bitrate.

Our simulation analysis thus shows the feasibility for our pre-response decoder to be used in a BCI framework in order to improve task performance specifically when the task is difficult and errors are costly.

## 4. Discussion

In this work, we investigated the neural correlates of confidence in a task with realistic stimuli, showed that these correlates are stimulus-locked rather than response-locked using a novel experimental design, found that confidence can be reliably decoded before a decision is made and without help from response-related activity, and developed a simulated BCI framework to show how and in what task conditions decoded confidence can be used to improve performance on a stimulus discrimination task. We incorporated a sufficiently long gap between stimulus and response in our task to allow stimulus- and response-locked brain activity to fully separate. Given this task design, our ERP and source localization analyses revealed that confidence-related activity is stimulus-locked. In the absence of a gap, however, this same activity appeared in the response-locked epoch due to leakage of stimulus-locked activity. Using this gap task, we then showed that purely using stimulus-locked pre-response activity, we could decode confidence and do so better with an SVM classifier compared to a battery of other classifiers. Finally, we used a simulated BCI framework to show that a classifier with this level of performance observed in real EEG data can indeed be used in a BCI in order to improve task performance especially for difficult high-stakes tasks.

### 4.1 The importance of a post-stimulus gap

We showed that the presence of a sufficiently long post-stimulus gap impacts whether or not confidence-related activity is observed to be stimulus-locked or response-locked. This observation was enabled by our new experimental design that carefully picked the gap length and consisted of two task conditions with the only difference being this gap. In particular, a comparison of neural activity between two stimulus discrimination tasks that differ only in terms of the presence of a gap had remained largely unexplored. This experimental design allowed us to conclude that confidence related activity is stimulus-locked and that confidence can be decoded purely from pre-response activity without any help from response-related activity.

As mentioned above, one reason that a post-stimulus gap is important is that it prevents stimulus- and response-locked activity from overlapping. This is, however, not the only reason a post-stimulus gap should be considered. For example, it has been shown that an ERP known as the error negativity (ERN) occurs in the response-locked epoch when stimulus processing continues after an incorrect response is made [94]. ERN amplitudes are typically stronger when a subject is aware that they made an error. It is possible that a long post-stimulus gap prevents subjects from making hasty decisions that lead to ‘avoidable’ errors that they are aware of, thereby attenuating the ERN. This phenomenon could possibly explain the lack of a response-locked difference between confident and unconfident conditions in the gap task. In studies where confidence, but not error monitoring, is the cognitive state of interest, a gap may be necessary to dissociate confidence-related activity from error-related activity.

Another phenomenon to consider is that of *temporal scaling*. Prior work has shown that rather than occurring at fixed latencies relative to events, certain neural responses can stretch or compress to fill a stimulus-response gap [95–97]. When characterizing particular neural responses, it is important to understand whether the response occurs at a fixed latency, or if it scales with the stimulus-response gap. Among other reasons, this distinction is important because fixed-latency responses may appear in the response-locked epoch with a small gap, while temporally-scaled responses may not. One way of making this distinction is to compare activity between tasks where the only difference is an enforced stimulus-response gap. In our study, we showed that in the absence of a gap, stimulus-locked and response-locked sources of neural activity do indeed overlap, which can suggest that the neural activity we observed occurs at a fixed latency relative to stimulus.

### 4.2 Confidence classifier is viable for use in a real-time BCI

The use of the gap task allowed us to explore whether, using just stimulus-locked pre-response activity, we can decode confidence. Indeed, the gap duration was chosen to be long enough to prevent response-related activity from interfering and confounding the decoding results. Our confidence classification analysis on this gap task showed that various classification algorithms could classify confidence from single-trial stimulus-locked pre-response EEG activity alone. Interestingly, among the considered classifiers, SVM achieved the highest accuracy, even outperforming the more sophisticated deep learning methods. In order to prevent overfitting on our dataset, we had to keep the size and number of layers in our MLP and CNN classifiers relatively small. It is likely that neural network models could use more complex architectures and achieve higher performance if more data is collected from each subject. However, collecting more training data would come at the cost of time and comfort for the subjects and future users of a BCI, and as such may not be desirable. Here, with a manageable number of trials (640) per subject, we were able to achieve a relatively high AUC of .76.

Beyond the classifier performance itself, an important point of consideration is whether this classifier is viable for use within a BCI framework. Viability requires not only that the classifier’s AUC is sufficient to allow for the BCI to improve the user’s performance, but also that the classifier can be run in real time. In section 3.4, we performed an extensive simulation that verified the first point, that our classifiers were accurate enough to enable a BCI for decision making. Regarding the second point, we note that all the classifiers we built, once trained, can classify single trials almost instantaneously. For these reasons, these confidence classifiers are indeed viable for use in a real-time BCI.

### 4.3 BCI Applications

We have shown that within our simulation framework, a realistic, imperfect confidence decoder can be used to improve performance on a stimulus discrimination task. This simulation assumes that the BCI has the ability to repeat the stimulus, i.e., that the task takes place in a controlled environment where the stimulus presentation can be controlled by a computer. Further, since the decoder uses stimulus-locked data to estimate the user’s confidence, it must have access to the stimulus onset times. It is important to consider whether it is reasonable to expect these conditions to hold in realistic use cases, and what can be done in cases where they do not.

In some realistic situations, it is entirely possible that the stimulus can be controlled by the BCI. For example, in assembly line quality control, a human inspector must look at samples pulled from the assembly line and determine whether or not they are up to standard. We envision a use case where the inspector is presented with images of products coming off the assembly line and must determine whether they are defective or not. In such a case, our BCI setup is directly applicable, as it is possible for the BCI to repeat or adjust the presentation of such images. Additionally, since the BCI can control when the images appear, the decoder would have access to stimulus onset times.

There are, however, situations where the aforementioned conditions do not hold. If the user’s task involves reacting to and making decisions regarding stimuli that occur naturally, at random, or in some other way that cannot be controlled by a computer, then our BCI framework would require some adjustments to be applicable. First, an event-detection method is needed so that stimulus-locked data can be collected. Prior work has developed algorithms that can detect behavioral or stimulus events in real time from neural activity [98,99]. Such algorithms can be used alongside the BCI in order to determine when stimuli appear. Once a stimulus is detected, neural data following the stimulus can then be fed into a classifier as usual. Second, if the BCI cannot repeat the stimulus, then some other feedback method is required in order to improve the user’s decision performance. Possible alternatives to stimulus repetition include providing additional task-relevant information to the user [42], providing input from AI or human agents performing the same task, or simply informing the user that they are unconfident and should take more time before making a decision. With the use of an event detection algorithm and an appropriate feedback signal, it should be possible for a BCI to improve the user’s performance even when stimulus times are unknown and cannot be controlled by a computer.

### 4.4 Future directions

We have shown with numerical simulations that our decoder can be used to improve task performance in a BCI framework. Specifically, the BCI improves performance when the cost of error is high, and the task is difficult. An important future direction would be to implement and test such a BCI with human subjects. The end goal of this research would be to develop a BCI that can assist human users in practical, real-world tasks. As stepping stones to this end goal, BCIs can be constructed for and tested with increasingly complex and more realistic tasks. One such complexity would be a task in which stimuli cannot be controlled by the BCI, as discussed in section 4.3. This would require the use of event-detection algorithms [98,99], and the development and testing of feedback methods that do not involve controlling the stimulus itself as discussed in section 4.3. Future work can explore the use of dynamical modeling methods in the development of decoding algorithms to improve cognitive state decoding [100–108]. Further, adaptively tracking the changes in EEG signals over time or detecting switches in this activity can further improve the performance of such BCIs [109–116].

**Figure A1:**
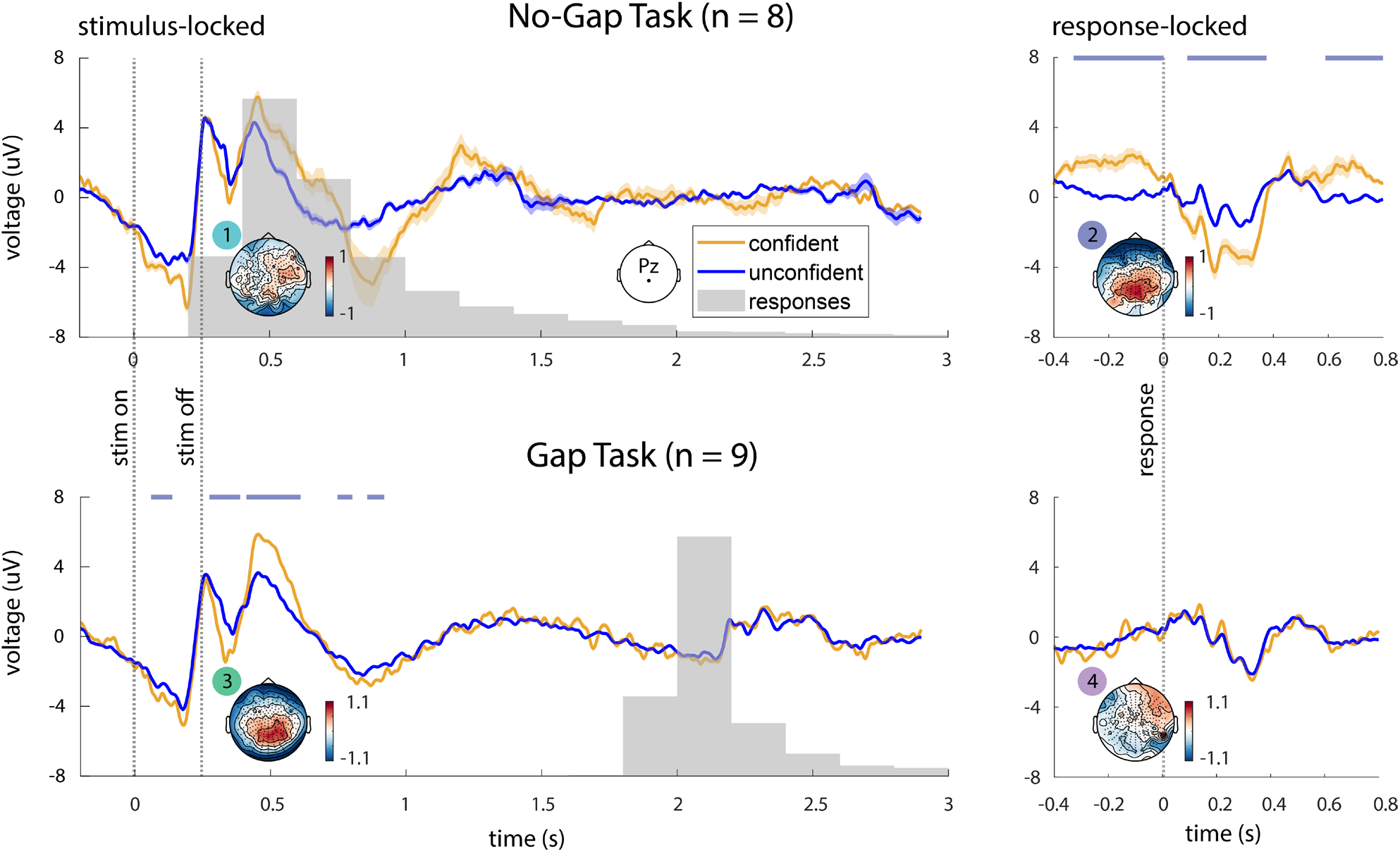
ERP Analysis using all trials. We repeated the ERP analysis shown in Figure 4, but without excluding any trials. Instead, all trials at and above the 80^th^ percentile of confidence reports were placed in the ‘confident’ group, while the remaining trials below the 80^th^ percentile were placed in the ‘unconfident’ group (in Figure 4, we again considered the trials above 80^th^ percentile as confident but only considered trials below 20^th^ percentile as unconfident unlike here). Results are qualitatively similar to the original analysis – the stimulus- and response-locked epochs of the no-gap task have differences over parietal channels, but when a gap is added, this pattern is revealed to be stimulus-locked.

## Appendix A

We now briefly explain the procedure for the cluster-based permutation test [57]. We consider EEG data from a single channel for two experimental conditions, c1 and c2. For the purpose of this explanation, each trial is s samples long and has a label corresponding to its experimental condition. To perform the cluster-based permutation test, we first compute t-values at each of the s samples to compare conditions c1 and c2. We then select all samples with t-values above a specified threshold. For our analysis, t-values were thresholded above the 99^th^ percentile. After thresholding, selected samples that are temporally adjacent are grouped into clusters. In other words, if two samples are both above threshold and temporally adjacent, they will be part of the same cluster. For each cluster, we compute a cluster statistic by taking the sum of t-values for all samples within the cluster. This clustering procedure (all steps until this point) is then performed for 1000 random permutations of the trial labels. For each of these permutations, the maximum cluster statistic is recorded. For each cluster that was computed using the true labels, the cluster-level p-value is the proportion of permutations where the maximum cluster statistic is greater than that cluster’s statistic, i.e. the sum of t-values for all samples within the cluster. Clusters were declared to be significant if their cluster-level p-values were less than .01.

## Appendix B

We repeated the ERP analysis described in Section 2.4.1 without excluding trials with confidence reports below the 80^th^ percentile and above the 20^th^ percentile (Figure A1). In this analysis, the previously excluded trials were incorporated into the ‘unconfident’ group. The results are largely similar to the main analysis – a large difference between conditions is seen in the response-locked epoch of the no-gap task, but in the gap task, this difference is instead stimulus locked. In the main analysis (Figure 4), we chose to exclude the middle 60% of trials in order to accentuate the difference between confident and unconfident conditions.

**Figure A2:**
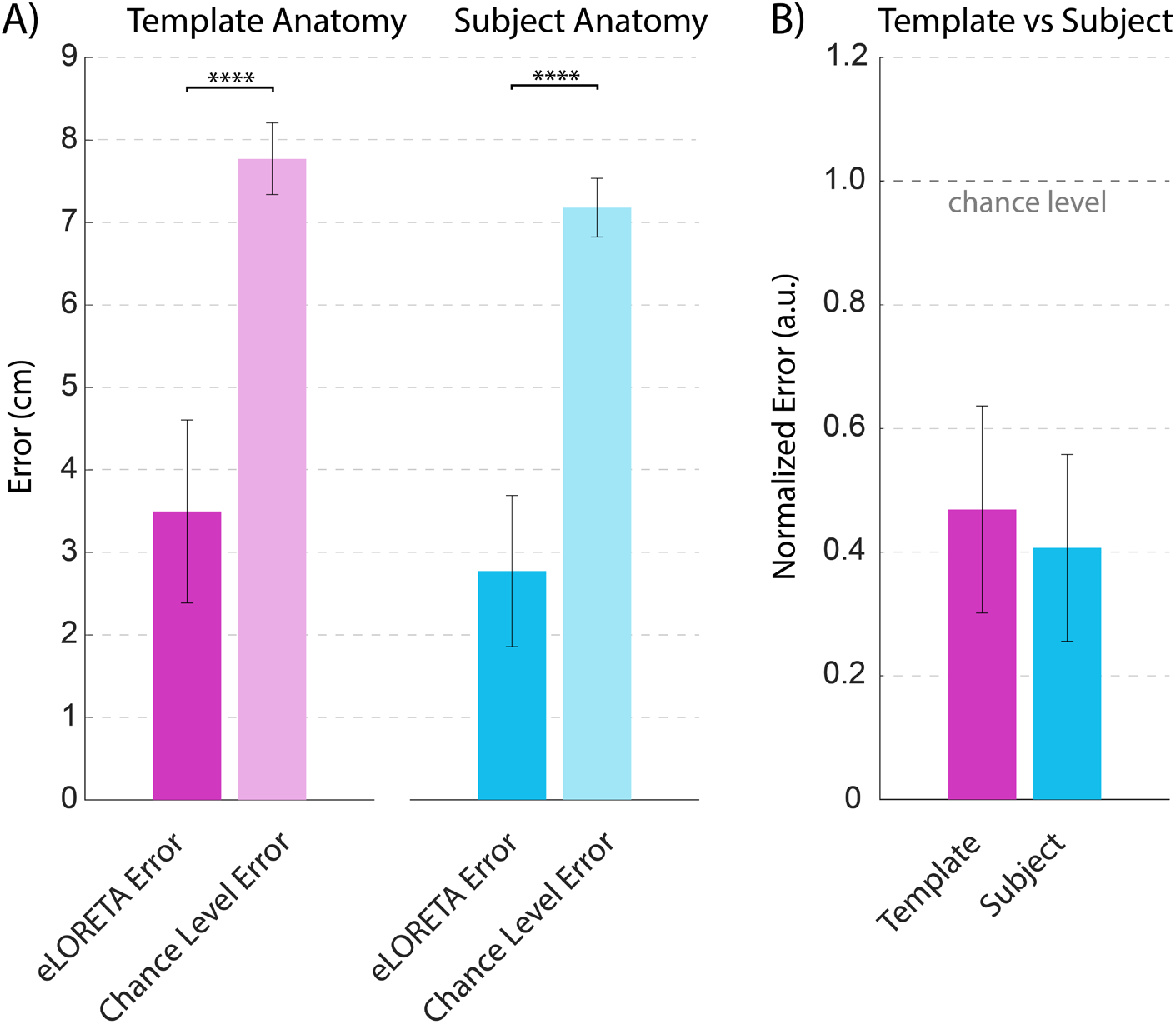
eLORETA performance comparison for template vs subject-specific anatomical models. (A) Comparison to chance level. For both anatomy types, eLORETA performed significantly better than chance (template anatomy p = 1.2e-12, subject anatomy p = 7.9e-15, paired t-test). (B) Comparison between template and subject anatomy, normalized by chance level. Although normalized eLORETA error with subject anatomy was slightly less than with template anatomy, this difference was not significant (p=.1).

## Appendix C

We used data from the publicly available Localize-MI dataset in order to compare the performance of eLORETA source localization while using subject-specific anatomy vs template anatomy [65]. The dataset features simultaneous EEG and intracranial stimulation, with the stimulation electrode positions serving as the ground truth for source localization analysis. Specifically, 7 subjects participated in 5-10 neural recording sessions with stimulation. Within each session, the stimulating electrode was kept at a fixed location, and 40-60 stimulation trials were performed, with one stimulation pulse per trial. The locations of the stimulating electrodes serve as the ground truth for source-localization methods and allow for their evaluation. This dataset also contained head models based on subject MRIs, and therefore allowed us to assess our eLORETA analysis pipeline with both subject-specific anatomy and template anatomy.

We performed this assessment by comparing the localization error of our method with a chance-level error to determine if the localization error was significantly better than chance. We applied eLORETA to trial-averaged EEG data from each session. The localization error was computed as the distance between the stimulating electrode and the source voxel with the highest current density. We computed the chance level error by taking the average distance from the stimulating electrode to every voxel in the eLORETA source space. This represents the expected error if the location of maximum current density was chosen as a random voxel with a uniform probability distribution. We then compared the performance of template and subject anatomy after normalizing each session’s error by its chance level. This was done to account for the difference in chance level between anatomy types.

Our analysis revealed that eLORETA with both subject-specific and template anatomy performed significantly better than chance level (Figure A2-A). The results using subject anatomy were not significantly better than results using template anatomy (Figure A2-B). These results validate our source localization pipeline and indicate that template anatomy can indeed be used to perform reliable source analysis.

## Acknowledgements

We thank Han-Lin Hsieh, Dongkyu Kim, Eray Erturk, Lucine Oganesian, Alireza Ziabari, Parsa Vahidi, Rahul Nair, Christian Song, Mustafa Avcu, and Christoph Schneider in the Shanechi Lab for their assistance in preparing subjects and collecting EEG Data. We thank Rahul Nair for his early contributions to the analysis code. We thank Riccardo Poli from the University of Essex for providing images and 3D models that were used in the experiments. This work was supported in part by the Army Research Office (ARO) under contract W911NF1810434 under the Bilateral Academic Research Initiative (BARI).

## Author Contributions

NS, OS, MS conceptualized the project and experimental design. NS, OS, and PA performed data collection. NS performed the analysis. MS supervised the project.

